# STED microscopy reveals mitotic stage-dependent CENP-A organization

**DOI:** 10.64898/2026.09.03.748884

**Authors:** Thomas C.Q. Burgers, Sietse J. Dijt, Luca Vitrano, Geert van den Boogaart, Rifka Vlijm

**Affiliations:** Molecular Biophysics, Zernike Institute for Advanced Materials, University of Groningen, Nijenborgh 7, 9747 AG Groningen, The Netherlands; Molecular Immunology, Groningen Biomolecular Science and Biotechnology Institute, University of Groningen, Nijenborgh 7, 9747 AG Groningen, The Netherlands

## Abstract

Chromosome segregation is vital. Its disruption can cause aneuploidy, a hallmark of cancer. Centromere protein A (CENP-A) is an important protein during the segregation as it marks the location of the centromere, where the kinetochore assembles for microtubule attachment. Each mammalian centromere contains hundreds of CENP-A nucleosomes, whose spatial arrangement is expected to be critical for kinetochore function and error-free chromosome segregation. However, previous studies— mostly in fixed cells and focused on metaphase—have yielded conflicting results on CENP-A organization. Given the centromere’s sub-diffraction size, we used stimulated emission depletion (STED) super-resolution microscopy to visualize the CENP-A organization in both living (about 45 nm resolution) and fixed (about 30 nm resolution) immortalized human retinal pigment epithelial cells (hTERT RPE-1). We found that CENP-A organization does not adopt a single architecture but spans a spectrum from dense clusters to fragmented subclusters, with mitotic stage-dependent abundance and morphology. CENP-A chromatin is most dispersed in prophase, and progressively compacts during prometaphase and metaphase as microtubules attach. In meta- and anaphase predominantly dense organizations are formed, often with a plate-like morphology. Yet, non-dense and non-plate-like organizations persist through metaphase and anaphase, suggesting CENP-A spatial reorganization is heterogeneous during cell division.

## Introduction

Cell division is essential for the persistence of life, requiring precise separation of the sister chromatids during mitosis. The physical segregation of chromosomes occurs at the centromere. Here, the DNA not only contains canonical nucleosomes for packaging, but also nucleosomes in which histone H3 is replaced by centromere protein A (CENP-A), marking centromeric chromatin (1). These CENP-A-containing nucleosomes define the sites for kinetochore assembly, giving rise to a multi protein complex that mediates microtubule attachment and force transmission during mitosis (2–6). Defects in kinetochore assembly can lead to genome instability such as aneuploidy, a hallmark of cancer (7), highlighting the importance of understanding how CENP-A organization supports accurate chromosome segregation.

Despite its central role, the spatial organization of CENP-A at human centromeres remains debated. In contrast to budding yeast, where a single CENP-A^Cse4^-containing nucleosome defines each centromere (8), human centromeres contain approximately 100 CENP-A nucleosomes during mitosis—measured on retinal pigment epithelial RPE-1 cells (9)—positioned together within a diffraction limited volume of about 200 to 300 nm (10, 11). This dense packing is important for centromere identity (12) and suggests an underlying higher order structure; however, no consensus has been reached regarding its organization. Most structural insights into centromeric chromatin have been derived from chemically fixed and often synchronized cells, with a predominant focus on metaphase cells or isolated mitotic chromosomes. As the centromere has a size that is comparable to the diffraction limit, imaging its arrangement requires approaches that are able to surpass the diffraction limit (∼250 nm). Super-resolution microscopy, expansion microscopy, and electron microscopy studies have reported various CENP-A arrangements, including compact clusters (13), ring-like distributions (14) and bipartite configurations (15). A study using super-resolution stochastic optical reconstruction microscopy (STORM) reported that in metaphase CENP-A typically forms a compact domain of about 235 nm (13). Using centromere chromosome orientation fluorescence in situ hybridization (Cen-CO-FISH) combined with structured illumination microscopy (SIM), alpha satellite DNA was found to adopt a ring-like arrangement, with a central dip in a subset of centromeres (14). The same study also used expansion microscopy in colcemid-arrested RPE-1 cells, showing heterogeneous crescent-like CENP-A patterns of about 150 to 200 nm (14). In addition, a recent study combining expansion microscopy and SIM to perform high-resolution imaging of unsynchronized human cells found bipartite centromere arrangements (15). In another study, imaging with STORM and MINFLUX showed that CENP-A forms a cluster of about 300 nm at each sister centromere containing smaller subclusters, with cluster size decreasing upon CENP-C depletion (16). These observations are consistent with cryo-electron tomography (cryo-ET), which revealed centromeres in DLD-1 cells to contain on average 30 nucleosome-associated complexes of about 20 to 25 nm (17). Overall, these studies report varied CENP-A organizations in mitosis with different interpretations of the underlying structure, with some resolving multiple small subclusters and others predominantly observing two domains at metaphase. These differences may arise from resolution limitations or variations in sample preparation. Moreover, it is still unclear how CENP-A organization evolves throughout mitosis.

Here, we use stimulated emission depletion (STED) super-resolution microscopy to resolve the CENP-A organization. By performing both fixed- and live-cell imaging, using varying labelling strategies (immunofluorescence and HaloTag (18) tagging of endogenous *CENPA*), across all mitotic stages we aim to provide a comprehensive, yet minimally perturbed view of centromeric chromatin organization during chromosome segregation. We observe that CENP-A is organized in dense clusters, fragmented subclusters, or an intermediate configuration, but not prominently as bipartite distribution. In addition, we identify mitotic stage-dependent differences in CENP-A organization, with more fragmented configurations at the start and end of mitosis and more compact arrangements during metaphase and anaphase. These observations reveal that CENP-A organization is heterogeneous and dynamically rearranged throughout mitosis. Together, this provides a framework to reassess current models of centromere organization and inheritance.

## Results

STED imaging resolved the spatial organization of CENP-A at mitotic centromeres into discrete nanoscale clusters, which exhibited a striking heterogeneity across all mitotic stages (Fig. 1). To systematically define these architectures across mitosis, we imaged immunolabelled CENP-A in unsynchronized, fixed, immortalized human retinal pigment epithelial cells (hTERT RPE-1) and assigned mitotic stages based on chromatin morphology visualized by confocal DAPI imaging (Fig. 1). Concurrent CENP-A and CENP-B imaging enabled identification of centromeres (Fig. 1), allowing consecutive STED imaging of the individual centromeres. The STED imaging of CENP-B allowed unambiguous discrimination between CENP-A signals derived from single versus closely adjacent centromeres.

**Figure 1.**
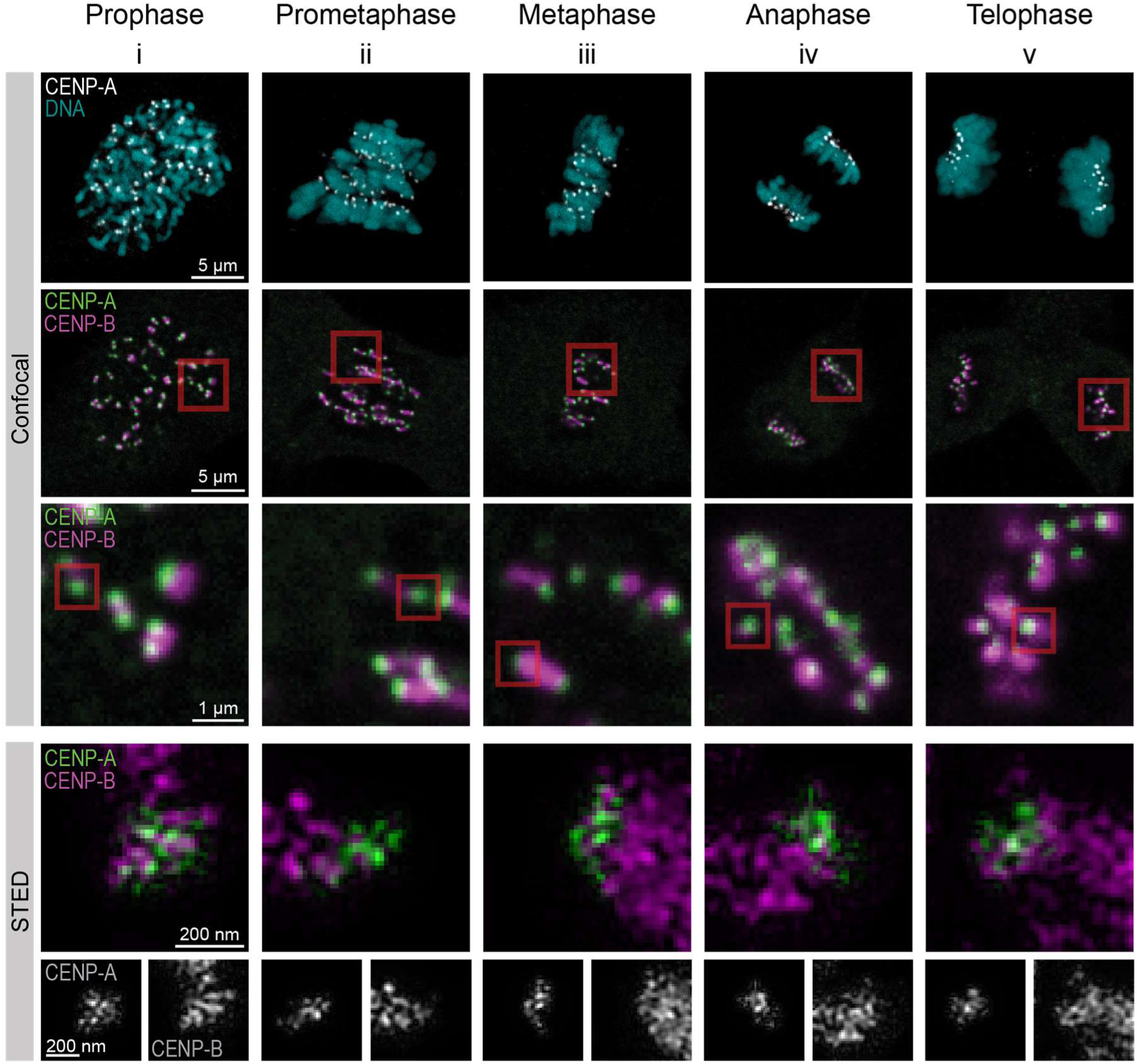
STED imaging of centromeres throughout mitosis in fixed hTERT RPE-1 cells. Representative confocal images of cells and STED images of centromeres in prophase (i), prometaphase (ii), metaphase (iii), anaphase (iv), and telophase (v). Cells were labelled with DAPI to show the DNA (cyan) and immunolabelled for CENP-A (green) and CENP-B (magenta). The red boxes indicate the areas magnified below. STED images were acquired using DyMIN and Wiener deconvoluted.

The substructures of CENP-A revealed by STED (Fig. 2A, Supplementary Fig. S1) showed broad heterogeneity, yet could be classified into three categories—dense, intermediate, or fragmented— based on the fragmentation within the centromere, the compactness of the CENP-A region, and the uniformity of the CENP-A distribution within the centromere. Scoring of all images (1019 centromeres from N = 3 replicates) revealed distinct trends across mitotic stages (Fig. 2A). Metaphase and anaphase showed the highest frequency of dense CENP-A arrangements (44-46%), while the intermediate class occurred at comparable frequencies across all stages except telophase, which displayed a slightly higher occurrence. The fragmented class was most prevalent in prophase and gradually decreased toward metaphase, followed by a modest increase toward telophase.

**Figure 2.**
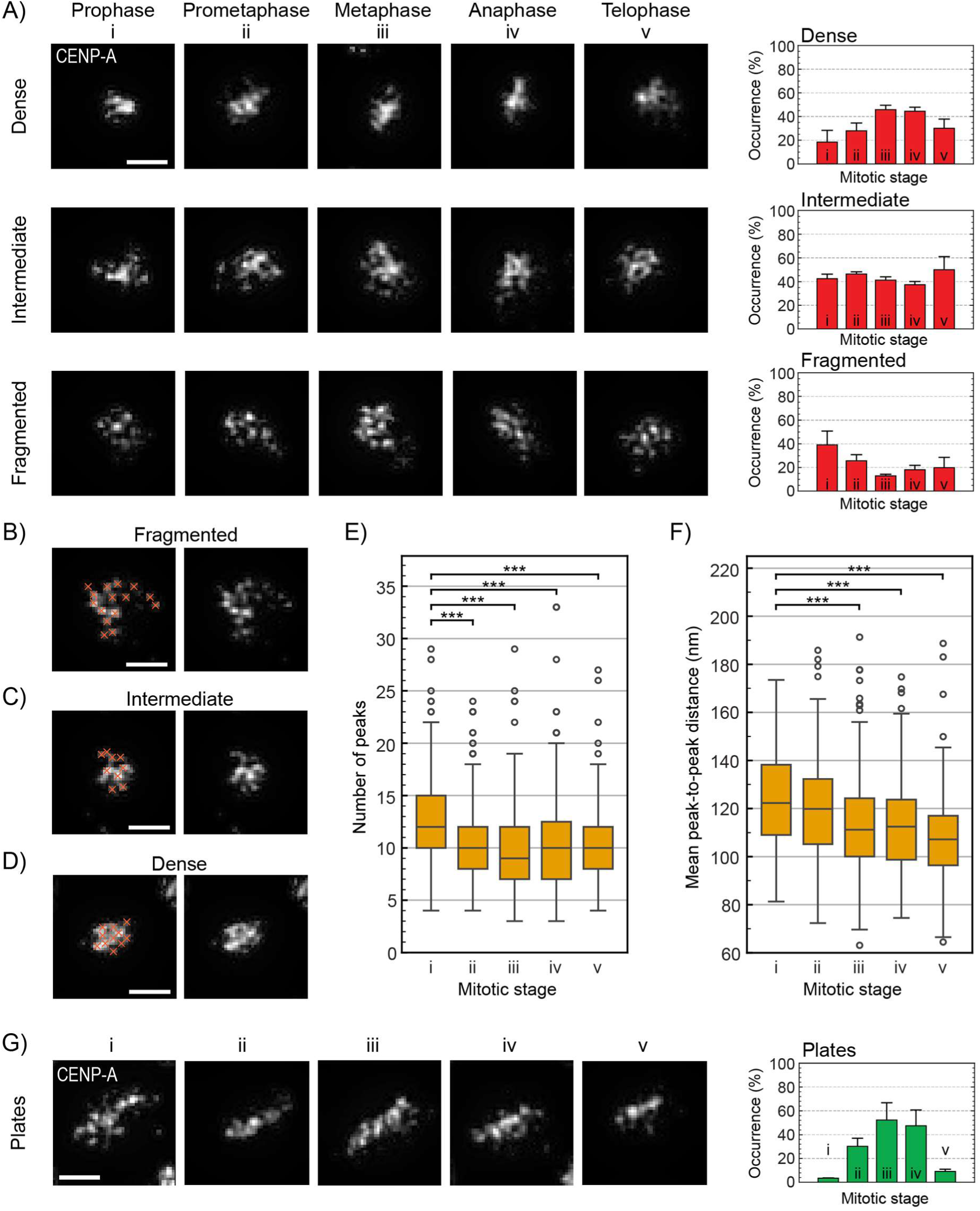
Analysis of CENP-A STED images in fixed hTERT RPE-1 cells. **A)** Examples of observed CENP-A organizations classified as dense, intermediate, or fragmented, for each mitotic stage (left, scale bar 200 nm) and the occurrences of each CENP-A organization per mitotic stage (right). **B-D)** Examples of a fragmented (B), intermediate (C), and dense (D) CENP-A organization showing the peaks (orange crosses, left images) found using local maximum peak fitting (scale bar 200 nm). **E)** Quantification of the number of peaks of all CENP-A organizations per mitotic stage. **F)** Quantification of the mean peak-to-peak distance per mitotic stage. Boxplots in E) and F) show the interquartile range, median center line, whiskers extending to the most extreme data points within 1.5x the interquartile range, and outliers. Significance was tested compared to prophase (i) using ANOVA followed by Dunnett’s multiple comparison test. Significant differences are shown using asterisks (*** = *p* < 0.001). Number of centromeres analysed per stage: N = 175 (i), 177 (ii), 328 (iii), 183 (iv), 151 (v) from 3 independent experiments). **G)** Examples of plate-like CENP-A organization, for each mitotic stage (left, scale bar 200 nm) and the occurrence of plates per mitotic stage (right). Bar charts in A) and G) show the mean ± SD of 3 independent experiments; number of cells/centromeres measured per stage: 11/175 (i), 12/178 (ii), 22/330 (iii), 12/184 (iv), 18/152 (v).

We applied image analysis algorithms to quantify the CENP-A organization in each centromere. To assess fragmentation, we used segmentation to identify separated CENP-A regions (Supplementary Fig. S2). The average number of CENP-A regions per centromere decreased from prophase (5.4 ± 2.7) toward metaphase (3.3 ± 1.8). In addition, we calculated the fragmented area, defined as the percentage of the area outside the largest region. This percentage is small when most CENP-A is in a single connected region and large when uniformly fragmented (19). The fragmented area decreased from prophase (20% ± 16%) toward metaphase (11% ± 13%), and was relatively low (15% ± 16% in all mitotic stages combined) compared to the average number of CENP-A regions (4.0 ± 2.4 in all mitotic stages combined), indicating that CENP-A is typically not uniformly fragmented.

To quantify nanoscale clustering, we used a peak-finding algorithm to identify individual clusters based on local intensity maxima (Fig. 2B-D). This analysis describes the clustered structure not only in clearly fragmented organizations, but also in denser CENP-A regions. The number of clusters decreased from prophase (13 ± 4.6) toward metaphase (10 ± 3.7), suggesting the structure becomes denser as clusters enlarge or merge due to proximity (Fig. 2E). The mean peak-to-peak distance per centromere slightly decreased from prophase (124 ± 20 nm) toward metaphase (113 ± 20 nm), indicating a less widely distributed structure (Fig. 2F).

To further analyse the CENP-A organization, we examined the shape of the CENP-A containing regions. We found that some regions adopted an elongated, plate-like organization (Fig. 2G, Supplementary Fig. S3). While these plate-like structures appeared in all mitotic stages, their frequency increased in prometaphase, peaking during metaphase and anaphase (Fig. 2G). Quantitative elongation measurements of all imaged structures confirmed this trend, as the average elongation of the overall CENP-A region was highest in metaphase and anaphase (Supplementary Fig. S4).

To detect and exclude fixation-induced artefacts, we used live-cell imaging as a complementary approach. While fixed-cell STED achieves superior resolution (∼30 nm, Supplementary Fig. S5), live-cell STED still achieves a significant resolution enhancement (∼45 nm, Supplementary Fig. S5) over conventional live-cell methods such as confocal imaging and SIM. We previously demonstrated the live-cell compatibility of STED microscopy (20), and used the same physiological conditions (controlled humidity, temperature, and CO₂) in this work. In hTERT RPE-1 cells with endogenous CENP-A-HaloTag (21), we visualized CENP-A-Halo using SiR-Halo. Our live-cell STED imaging of CENP-A (Fig. 3A, 645 centromeres from N = 3 replicates) recapitulated the observations made in fixed cells of the dense, fragmented and intermediate organizations (Fig. 3B), where prophase has a majority of fragmented organizations, with increasing occurrence of dense organizations towards metaphase. Analysis of the overall CENP-A morphology revealed plate-like organization, similar to fixed-cell imaging (Fig. 3C). Moreover, the trend in elongation was also similar to fixed-cell imaging, with the highest average elongation in metaphase and anaphase (Supplementary Fig. S4).

**Figure 3.**
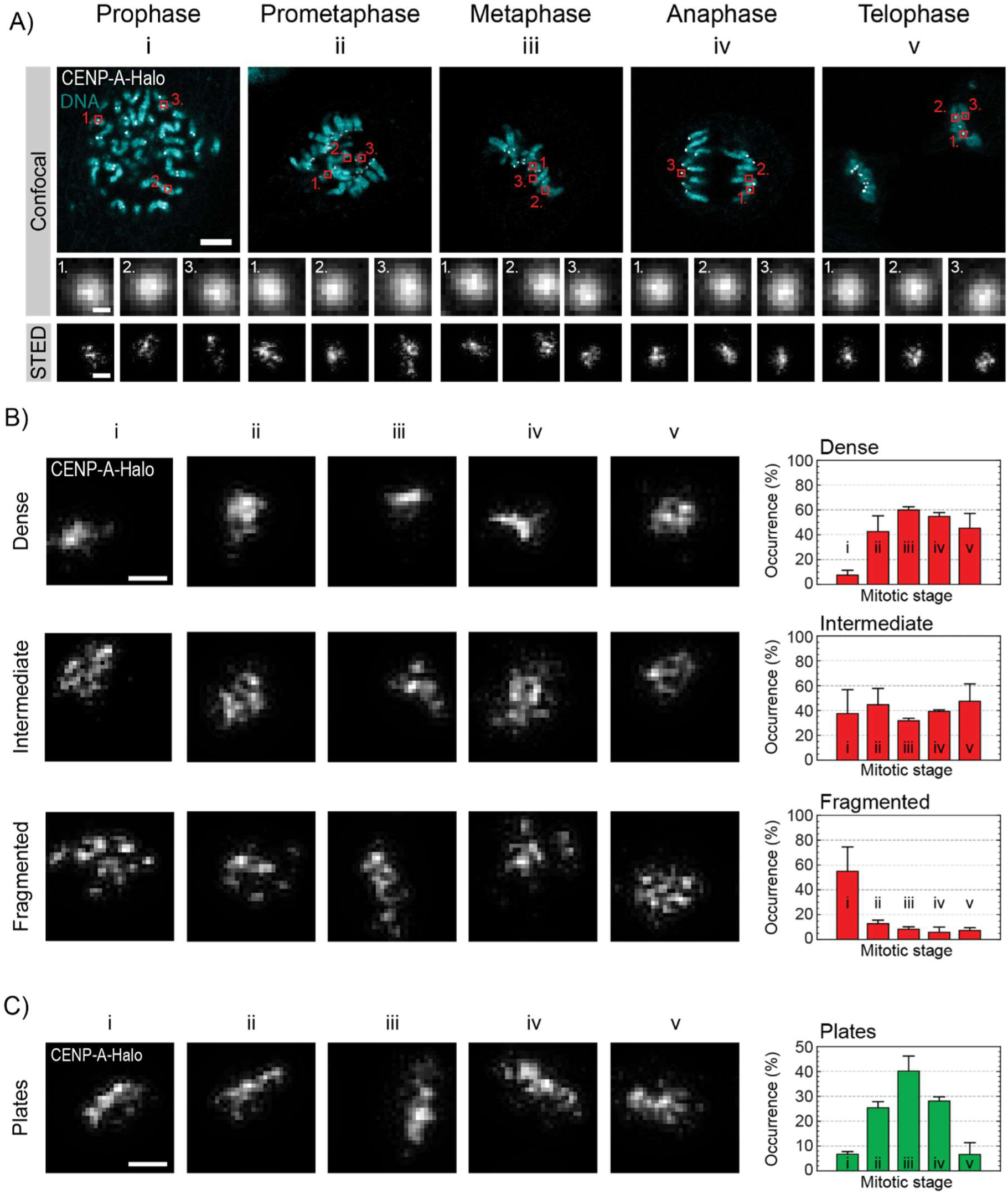
Live-cell STED imaging and analysis of CENP-A throughout mitosis. Live hTERT RPE-1 CENP-A-Halo cells were labelled with SiR-Halo and imaged under physiological conditions. STED images were acquired using DyMIN and deconvoluted using a Wiener filter for visualization. **A)** Representative overview images of cells in prophase (i), prometaphase (ii), metaphase (iii), anaphase (iv), and telophase (v) (top row, scale bar 2 µm). Below, confocal and STED images of three individual centromeres (indicative location marked by red squares with corresponding number in the overview) (middle and bottom rows, scale bar 200 nm). **B)** Representative dense, intermediate, and fragmented CENP-A spatial organization at mitotic centromeres observed in live-cell STED imaging (left, scale bar 200 nm) and the occurrences of each CENP-A organization per mitotic stage (right). **C)** Examples of elongated, plate-like CENP-A arrangements detected in live cells (left, scale bar 200 nm) and the occurrence of plates per mitotic stage (right). Bar charts in B) and C) show the mean ± SD of 3 independent experiments; number of cells/centromeres measured per stage: 9/95 (i), 13/139 (ii), 11/135 (iii), 11/114 (iv), 10/162 (v).

To probe the dynamics of CENP-A organization at the scale of seconds, we performed live-cell STED imaging of CENP-A-Halo, acquiring five consecutive frames at 1 Hz for cells across all mitotic stages (N = 105 centromeres). Within this timescale, we did not observe major changes in the shape and internal organization of CENP-A regions (Supplementary Fig. S6), indicating that the changes in CENP-A organization throughout mitosis in general occur at longer timescales. Due to the substantial photobleaching that is unavoidable for live-cell STED imaging at this resolution, and the high mobility of the individual centromeres which hinders tracking single centromeres over longer times, longer time-scales of the dynamical reorganization of CENP-A-Halo were not investigated.

## Discussion

In this work, we visualized the organization of CENP-A in hTERT RPE-1 cells throughout all mitotic stages, using live- and fixed-cell super-resolution microscopy in unsynchronized cells. We found for both sample preparation methods that CENP-A was organized heterogeneously, ranging from single dense clusters to fragmented subclusters. Throughout mitosis, we observed changes in the occurrence of these types of organizations and the shape of the CENP-A region, indicating dynamic rearrangement of CENP-A-containing DNA during mitosis. We did not find these rearrangements in CENP-A within short videos (1 Hz) of a few seconds, suggesting longer timescales for these rearrangements.

Our findings support and extend a model in which centromeres are composed of multiple smaller clusters of CENP-A containing nucleosomes. Averaged across mitosis, we detected 11 ± 4.2 clusters per centromere (∼9 nucleosomes per cluster, assuming ∼100 CENP-A nucleosomes per centromere (9)). Notably, out of all stages, prophase centromeres had on average the highest number of clusters. Our resolution sets a lower bound for cluster counts and an upper estimate for CENP-A nucleosomes per cluster. Especially the limited z-resolution of 2D STED (22) may merge height-separated clusters. With increasing compaction, such as upon transition from prophase towards metaphase, an increasing underestimation of the cluster count is expected. Given these constraints, our results align with previous STORM and MINFLUX CENP-A imaging (16) and cryo-ET observations of centromeres containing clustered nucleosome-associated particles which were hypothesized to be inner kinetochore complexes (17). Contrary to reports of stable bipartite metaphase architectures (15), our STED analysis of fixed and live hTERT RPE-1 cells revealed a spectrum of CENP-A spatial organizations (dense, intermediate, and fragmented), with metaphase centromeres exhibiting a preferred dense organization compared to the other mitotic stages.

The decrease in fragmentation of CENP-A organization after prophase, and change to a typically more elongated, dense arrangement in metaphase suggests that, as the kinetochore assembles towards metaphase, the underlying chromatin is also reorganized by bringing CENP-A chromatin in closer proximity. Although this trend was evident in both live and fixed cell imaging, dense, plate-like configurations are not the only found CENP-A organization in metaphase. Instead, all mitotic stages demonstrate variability in CENP-A chromatin organization. Since non-cancerous cells were used, under incubator conditions, without treatments such as synchronization, the presence of fragmented, non-plate-like CENP-A organizations in metaphase and anaphase, suggests that the organization of the CENP-A chromatin in a dense, plate-like configuration is not critical for successful cell division.

Interpretation of mitotic chromatin organization by fluorescence microscopy can be complicated by artefacts arising from fixation, labelling, or perturbations associated with cell-cycle synchronization. The combination of fixed- and live-cell imaging employed here mitigates several of these limitations. While the higher spatial resolution achieved in fixed cells enabled detailed characterization of CENP-A organization, detection of the same trends in live-cell imaging argues against fixation-induced artefacts. Likewise, the agreement between antibody-based labelling and endogenously HaloTag-labelled cells indicates that the observed organization is not specific to a single labelling strategy.

Previous studies have provided detailed views of centromere architecture at selected stages of mitosis, most prominently metaphase, and arrested cells. Here, imaging of asynchronous cell populations enabled systematic sampling across all major mitotic stages without synchronization. This broader stage coverage revealed pronounced changes in CENP-A organization during mitotic progression, and the consistency across imaging modalities supports the robustness of these stage-dependent observations.

Our results show that across mitosis, CENP-A organization does not predominantly adopt a specific architecture, such as a bipartite structure, but instead exhibits a continuum of states from dense clusters to fragmented subclusters. The abundance of these states and the overall CENP-A domain morphology vary by mitotic stage, indicating dynamic, rather than static, centromere chromatin organization. In prophase, CENP-A chromatin is most dispersed, potentially facilitating kinetochore assembly before microtubule capture, and progressively compacts during prometaphase and metaphase as microtubules attach. We hypothesize that dense, plate-like organizations in meta- and anaphase support stability under high pulling forces. These observations refine centromere models by revealing heterogeneous, stage-dependent organization. Yet, non-dense, non-plate-like organizations persist through metaphase and anaphase, suggesting spatial reorganization of CENP-A chromatin is not critical for cell division. These findings provide a framework for future super-resolution studies to resolve the spatial principles governing centromere organization.

## Methods

### Cell culture

hTERT RPE-1 cells (ATCC CRL-4000) (a kind gift from prof. dr. Marcel van Vugt, UMCG, Groningen) and hTERT RPE-1 CENP-A-Halo cells (a kind gift from prof. dr. Iain Cheeseman, Whitehead Intsitute, MIT, USA) (21) were cultured in Dulbecco’s Modified Eagle’s Medium (DMEM; ThermoFisher, 31966-021) supplemented with 10% fetal bovine serum (FBS; ThermoFisher, A5209502) and 1% penicillin/streptomycin (ThermoFisher, 15140122). Cells were grown at 37°C in humidified air with 5% CO_2_.

### Immunolabelling

hTERT RPE-1 cells were seeded on 18 mm No. 1.5H cover slips (Marienfeld Superior, 0117580) in a 12-well plate (TPP, 92012) and grown for 48-72 hours. Subsequently, cells were fixed with 4% formaldehyde (Sigma-Aldrich, 47608) diluted in PBS (Sigma-Aldrich, P2287) for 15 min at room temperature. After fixation, coverslips were washed three times with PBS.

Fixed cells were permeabilized using 0.2% Triton X-100 (Sigma-Aldrich, X100) in PBS for 15 min and washed three times with PBS. After blocking with blocking buffer (10% goat serum (ThermoFisher, 16210-072) in PBST (0.1% Tween20 (Sigma-Aldrich, P2287) in PBS)) for 1-2 hours at room temperature, cells were incubated for 1 hour with primary antibodies against human CENP-A (mouse IgG1, MBL, clone 3-19, D115-3MS, 1:100) and CENP-B (polyclonal rabbit IgG, Abcam, ab25734, 1:200) diluted in blocking buffer. The coverslips were washed four times with PBST and then incubated with secondary antibodies goat anti-mouse STAR RED (Abberior, STRED-1001) and goat anti-rabbit STAR 580 (Abberior, ST580-1002) diluted in blocking buffer. Both primary and secondary antibody labelling were performed by placing the coverslips on a 30 µL droplet of antibody solution on parafilm. After secondary antibody labelling, cells were washed twice with PBST, counterstained with 36 nM DAPI (GeneCopoeia) in PBST for 4 min and washed four times with PBST. Finally, the coverslips were mounted using 10 µL homemade Mowiol mounting medium (23).

### Live-cell sample preparation

hTERT RPE-1 CENP-A-Halo cells were seeded in culture dishes (CELLview, 627870) in FluoroBrite (ThermoFisher, A1896701) supplemented with 10% FBS, 1% penicillin/streptomycin and 4 mM GlutaMAX (ThermoFisher, 35050061). Cells were grown for at least 40 hours prior to labelling for 1 hour with 1 µM SiR-Halo (24) (a kind gift from dr. Alexey N. Butkevich and prof. dr. Stefan W. Hell, Max Planck Institute Göttingen, Germany) and 0.75 µM Hoechst 33258 (ThermoFisher, H3569) in 250 µL supplemented FluoroBrite. After labelling, cells were washed with supplemented FluoroBrite and finally placed in conditioned imaging medium (a 1:1 mixture of the seeding medium and fresh supplemented FluoroBrite).

### General STED microscopy

All measurements were performed on a STED microscope (Abberior Expert Line), equipped with an Okolab cage incubator, utilizing a 100x UPlanApo 100x/1.40 oil immersion objective (Olympus). A 0.1 µm TetraSpeck bead (Invitrogen, T7279) sample was used before each experiment to ensure correct laser alignment. The STED microscope was controlled through Imspector software.

A 40 MHz pulsed 775 nm STED laser (3 W at laser head) and a pulsed 640 nm laser (1.2 mW at laser head) were used for the excitation and depletion of SiR-Halo and STAR RED dyes. A pulsed 561 nm excitation laser (203 µW at laser head) and the 775 nm STED laser were used for excitation and depletion of the STAR 580 dye. DAPI and Hoechst 33258 were imaged only at confocal resolution using a continuous wave 405 nm excitation laser (20.7 mW at laser head).

To obtain the highest attainable resolution and/or number of frames, we used RESCue (25) and DyMIN (26) adaptive illumination strategies (Supplementary Fig. S7). Pixel illumination was stopped if the number of photons after a certain time is below a set threshold (i.e. background) or when the probe threshold is reached for probe masks (RESCue). We used probe masks at lower resolutions to gently probe the location of the structure and limit high-resolution STED imaging to the regions containing fluorophores (DyMIN). Together, these techniques prevent unnecessary photobleaching as fluorophores undergo fewer excitation and (stimulated) emission cycles within one image. Consequently, more images at the same resolution, or images at a higher resolution were obtained.

### Fixed-cell STED microscopy

STED microscopy of immunolabelled fixed cells was performed at room temperature. Mitotic stages were identified using confocal imaging of DNA. Simultaneous imaging of CENP-A and CENP-B allowed identification of individual centromeres. Consecutively, single centromeres were centred and positioned into focus based on the confocal CENP-A signal, and imaged at STED resolution. CENP-A STED was executed in a light-reducing DyMIN sequence containing three steps: a confocal probe image, an intermediate STED probe image, and a high-resolution STED image (Supplementary Fig. S7A). For the final high-resolution image, 6 frames were taken within the same probe mask to minimize spatially inhomogeneous bleaching while ensuring maximal signal. The final image was obtained by the sum of all 6 frames. The settings for the high-resolution CENP-A STED image were: 598 nm x 598 nm field of view with 13 nm pixels, 0.7 AU pinhole, 210 µs total pixel dwell time over 10 line repetitions, 1.1% 640 nm excitation power, 60% STED power, detection between 647-740 nm and a gating width of 9 ns.

### Live-cell STED microscopy

STED microscopy of living cells was performed with the incubator heated to 37°C (Okolab, H201-T-UNIT-BL), while supplying humidified 5% CO_2_ gas to the sample at 0.3 L/min flow rate by mixing pure CO_2_ (Westfalen) with air using a gas controller (Okolab, CO2-UNIT-BL), followed by passing a humidifier (Okolab, HM-VF). Mitotic cells were identified by Hoechst DNA labelling. CENP-A-Halo regions were selected, focused, and consecutively imaged at STED resolution. CENP-A-Halo was imaged in two steps: first a low resolution confocal probe image, followed by a high-resolution STED image. For single high-resolution STED images, we used one frame within the probe mask (Supplementary Fig. S7B). For time-lapses, the probe mask is measured in a separate line step in the same measurement as the high-resolution STED, such that the mask is updated for each frame (Supplementary Fig. S7C). The settings for high-resolution CENP-A STED images were: 640-888 nm square field of view with 20-24 nm pixels, 0.8 AU pinhole, 50-90 µs total dwell time over 1-2 line repetitions, 12-13% 640 nm excitation power, 24-53% STED power, detection between 647-740 nm and a gating width of 10 ns.

### STED image analysis

Images were processed using Wiener deconvolution (27). All raw data and analysis code is available on DataverseNL (28). Images were manually classified into three predefined categories (dense, partially fragmented, fragmented) by two independent researchers based on the fragmentation within the centromere, the compactness of the CENP-A region, and the uniformity of the CENP-A distribution within the centromere. Each researcher performed the classification separately and was blinded to the other’s assessments. Discrepancies were resolved through discussion until consensus was reached. From all datapoints an additional classification was performed (manually) to discriminate between a plate (elongated) or a non-plate structure based on the general shape of the overall CENP-A containing region, and ratio between length and width of the CENP-A region.

For the quantifications, single centromere masks were obtained by thresholding the raw STED images with a threshold of 20% of the maximum image intensity. Masks were manually corrected to include only a single centromere region if necessary. For the quantification of the number of CENP-A regions and fragmented area, we used a segmentation algorithm, which segmented Wiener-deconvolved images using a threshold of 27% of the maximum image intensity and distinguished regions using the *label* function with 1-connectivity from the scikit-image python package (29). The fragmented area was calculated as the percentage of the segmented area outside the largest region. For the quantification of CENP-A clusters, we employed a peak-finding algorithm, which determined all local maxima in the Wiener-deconvolved images within the single centromere mask and above 20% of the maximum image intensity through the *peak_local_max* function of scikit-image (29). For the quantification of elongation, the *regionprops* function of scikit-image (29) was used to obtain the major and minor axis lengths of a Legendre ellipse corresponding to the single centromere mask. The elongation was calculated as (30):

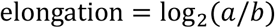

where *a* and *b* are the major and minor ellipse axis lengths, respectively (Supplementary Fig. S4A-B).

## Acknowledgements

We thank dr. Alexey N. Butkevich and prof. dr. Stefan W. Hell (Max Planck Institute for Medical Research, Heidelberg) for kindly providing the SiR-Halo dye. We thank prof. dr. Marcel van Vugt (University Medical Center Groningen, Groningen) for kindly providing the hTERT RPE-1 cell line. We thank prof. dr. Iain Cheeseman (Whitehead Intsitute, MIT, USA) for kindly providing the hTERT RPE-1 CENP-A-Halo cells.

## Author contributions

SJD, TCQB: contributed to experimental design, performing experiments, data analysis, writing. LV: contributed to preliminary experiments. GVDB: contributed to the conceptual framework, data discussion. RV: contributed to experimental design, data analysis, writing, and supervision. All authors contributed to reviewing and editing.

## Supplement

**Supplementary Figure S1.**
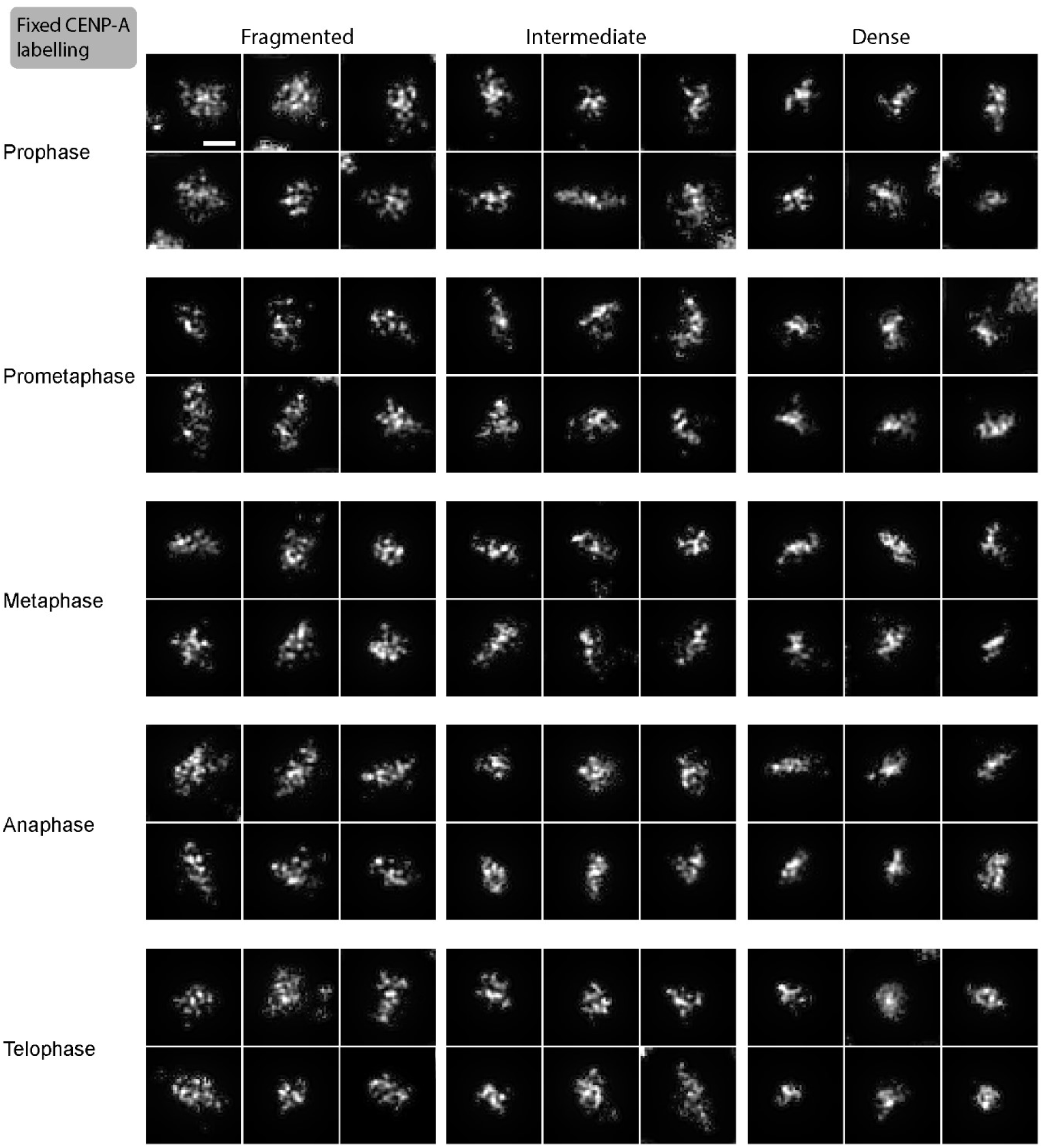
STED images of the spatial arrangement of all three classes of CENP-A organizations (fixed cells). Additional representative STED images of dense, intermediate, and fragmented CENP-A organizations across mitotic stages (Wiener filtered, scale bar 200 nm).

**Supplementary Figure S2.**
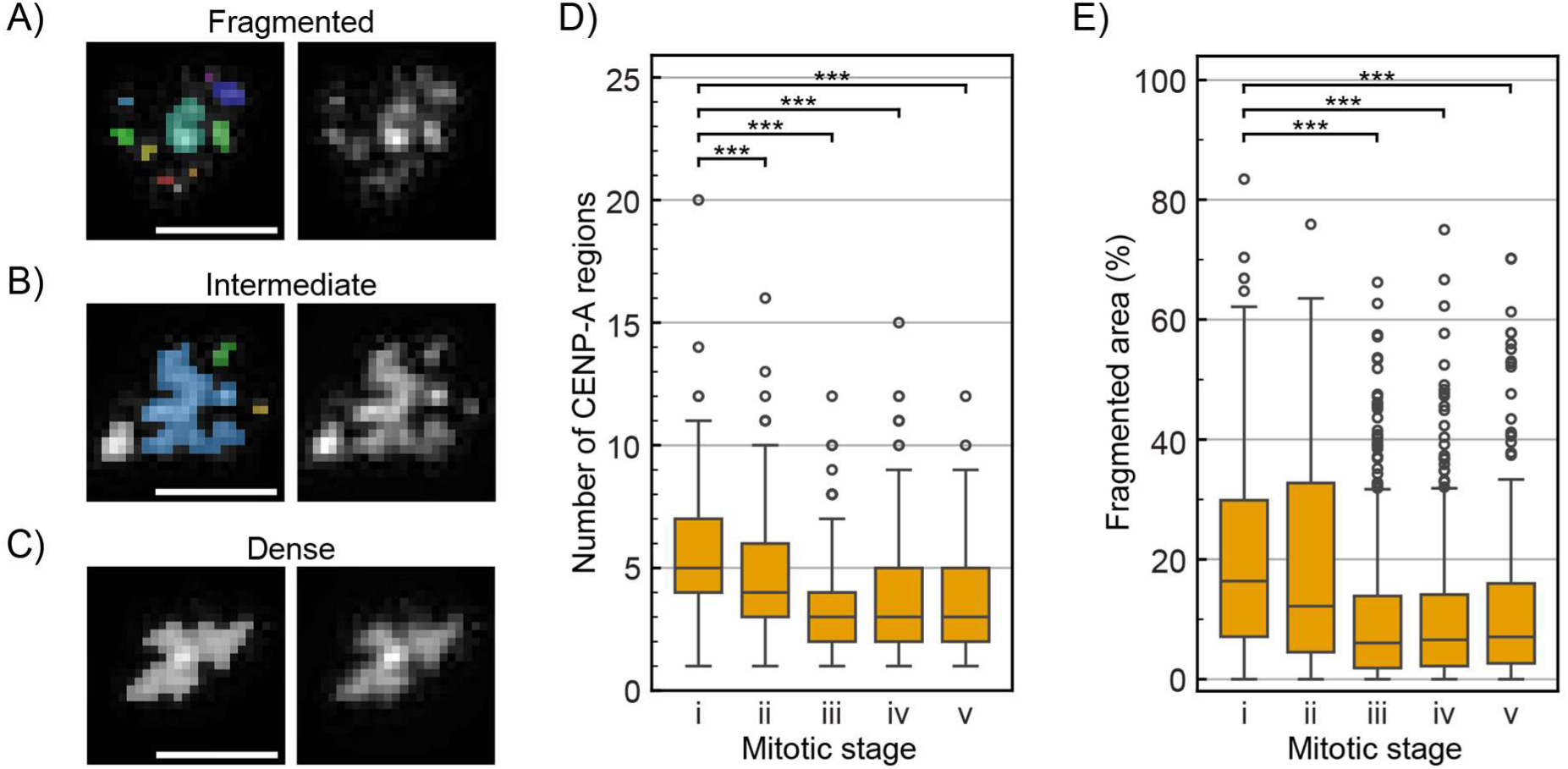
Fragmentation analysis of CENP-A STED images in fixed hTERT RPE-1 cells. **A-C)** Examples of a fragmented (A), intermediate (B), and dense (C) CENP-A organization showing the segmented CENP-A regions in different colors on the left image (scale bar 200 nm). **D)** Quantification of the number of CENP-A regions within individual centromeres per mitotic stage. **E)** Quantification of the fragmented area (percentage of the area outside the largest region) of individual centromeres. Boxplots in D) and E) show the interquartile range, median center line, whiskers extending to the most extreme data points within 1.5x the interquartile range, and outliers. Significance was tested compared to prophase (i) using ANOVA followed by Dunnett’s multiple comparison test. Significant differences are shown using asterisks (*** = *p* < 0.001). Number of centromeres analysed per stage: N = 175 (i), 177 (ii), 328 (iii), 183 (iv), 151 (v) from 3 independent experiments.

**Supplementary Figure S3.**
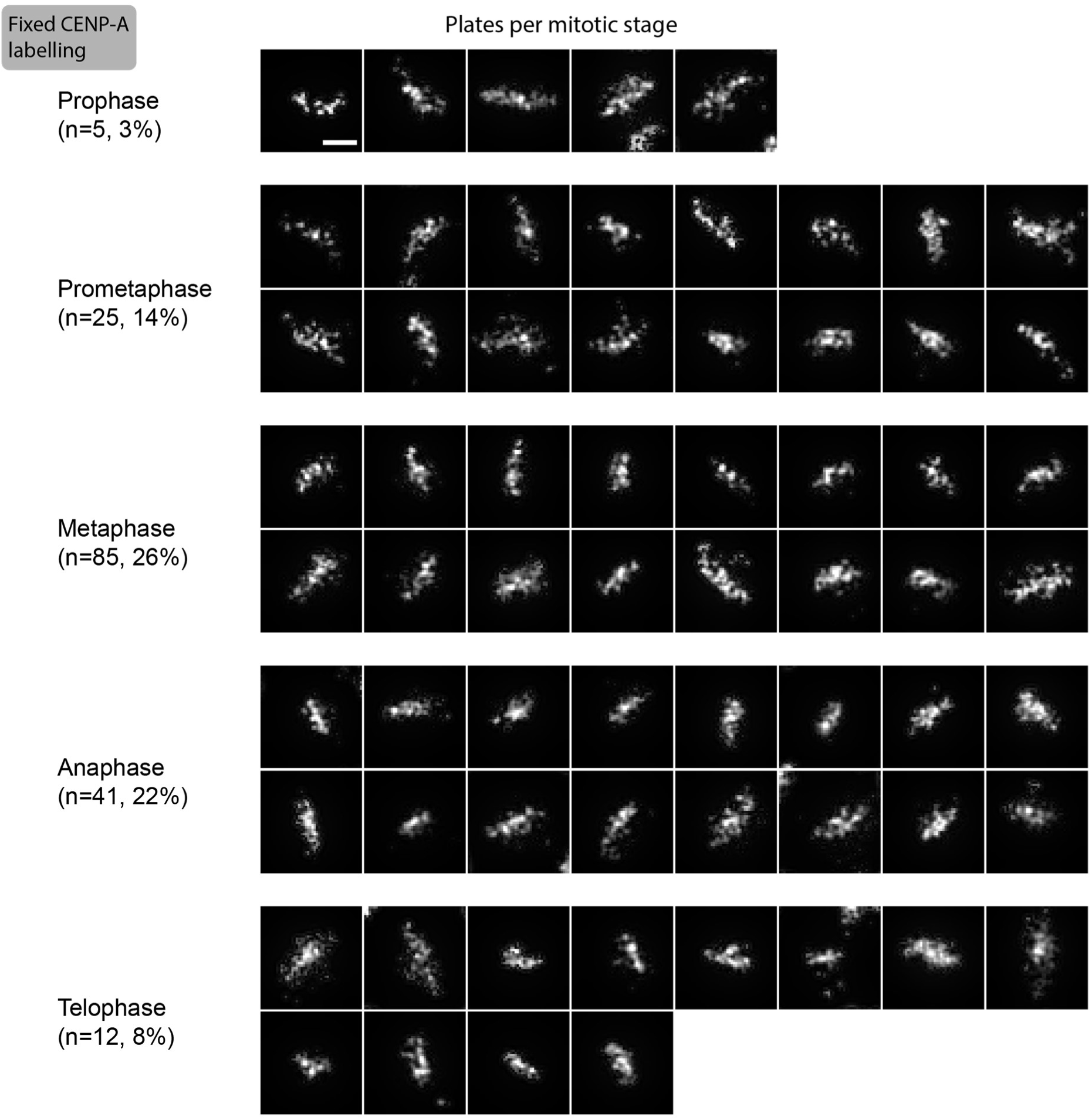
Plate-like CENP-A organization (fixed cells). Additional representative STED images showing elongated, plate-like CENP-A arrangements across mitotic stages. The number and percentage of observed plates per stage is indicated. (Wiener filtered, scale bar 200 nm).

**Supplementary Figure S4.**
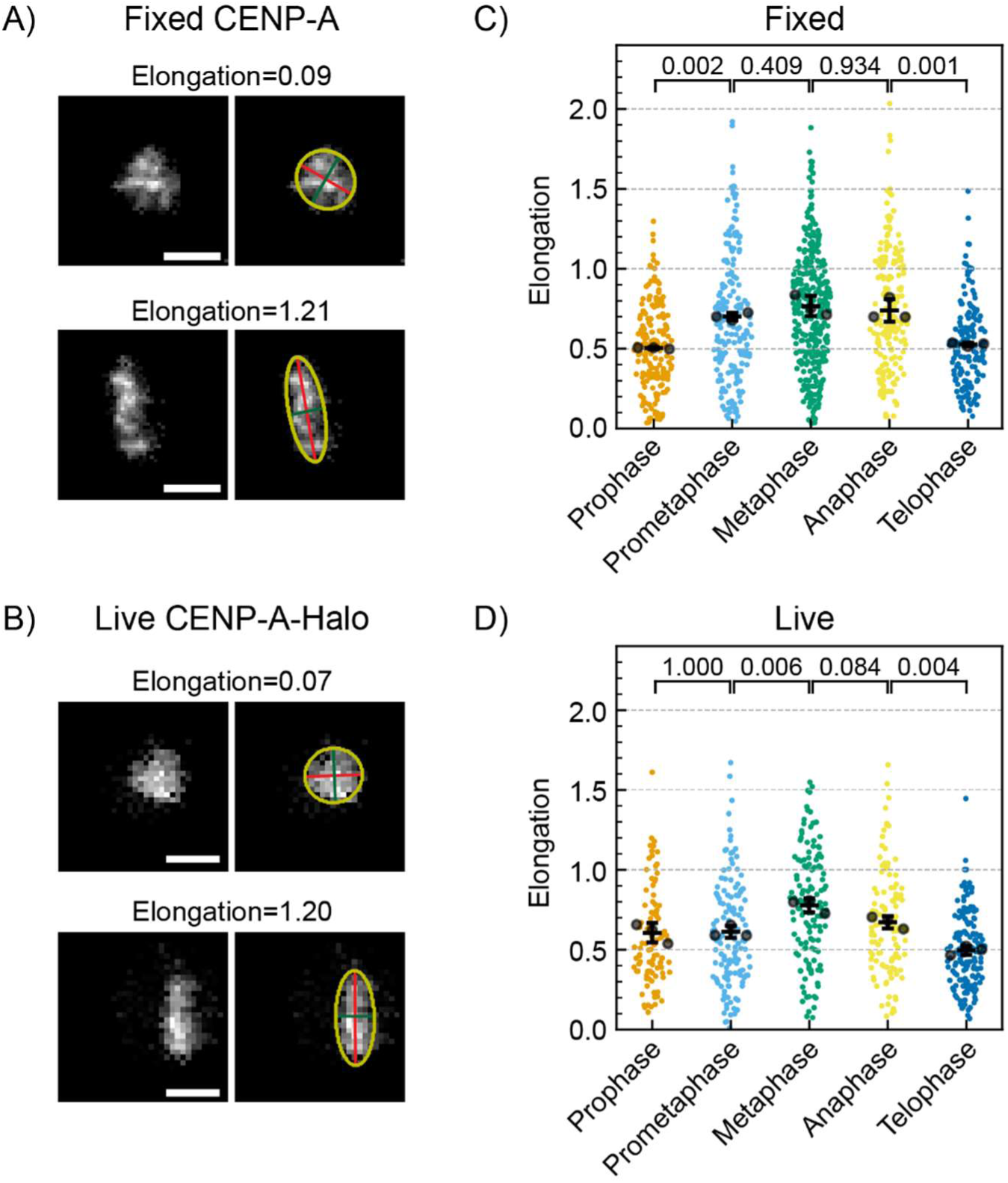
The elongation of the CENP-A region of individual centromeres throughout mitosis. The elongation of the overall CENP-A structure of individual centromeres was quantified from STED images of immunolabelled CENP-A in fixed cells (A/C) and CENP-A-Halo in live cells (B/D). Elongation is calculated as log_2_(*a*/*b*), where *a* and *b* are the major and minor ellipse axis lengths, respectively. An elongation of 0 indicates a circle and at an elongation of 1 the major axis is twice as long as the minor axis. **A/B)** Examples of STED images, showing the Legendre ellipse (yellow), major axis (red), and minor axis (green), with the corresponding elongation indicated above the image. Scale bars 200 nm. **C/D)** Quantification of elongation throughout mitosis, showing individual datapoints, the mean per replicate (large black circles) and mean ± SD of N = 3 independent experiments. Number of cells/centromeres: 11/176 (pro-), 12/177 (prometa-), 22/328 (meta-), 12/184 (ana-), 18/151 (telophase) for fixed and 9/93 (pro-), 13/139 (prometa-), 11/135 (meta-), 11/113 (ana-), 10/158 (telophase) for live cell imaging. *p*-values were calculated using Tukey’s test and are shown in the plot.

**Supplementary Figure S5.**
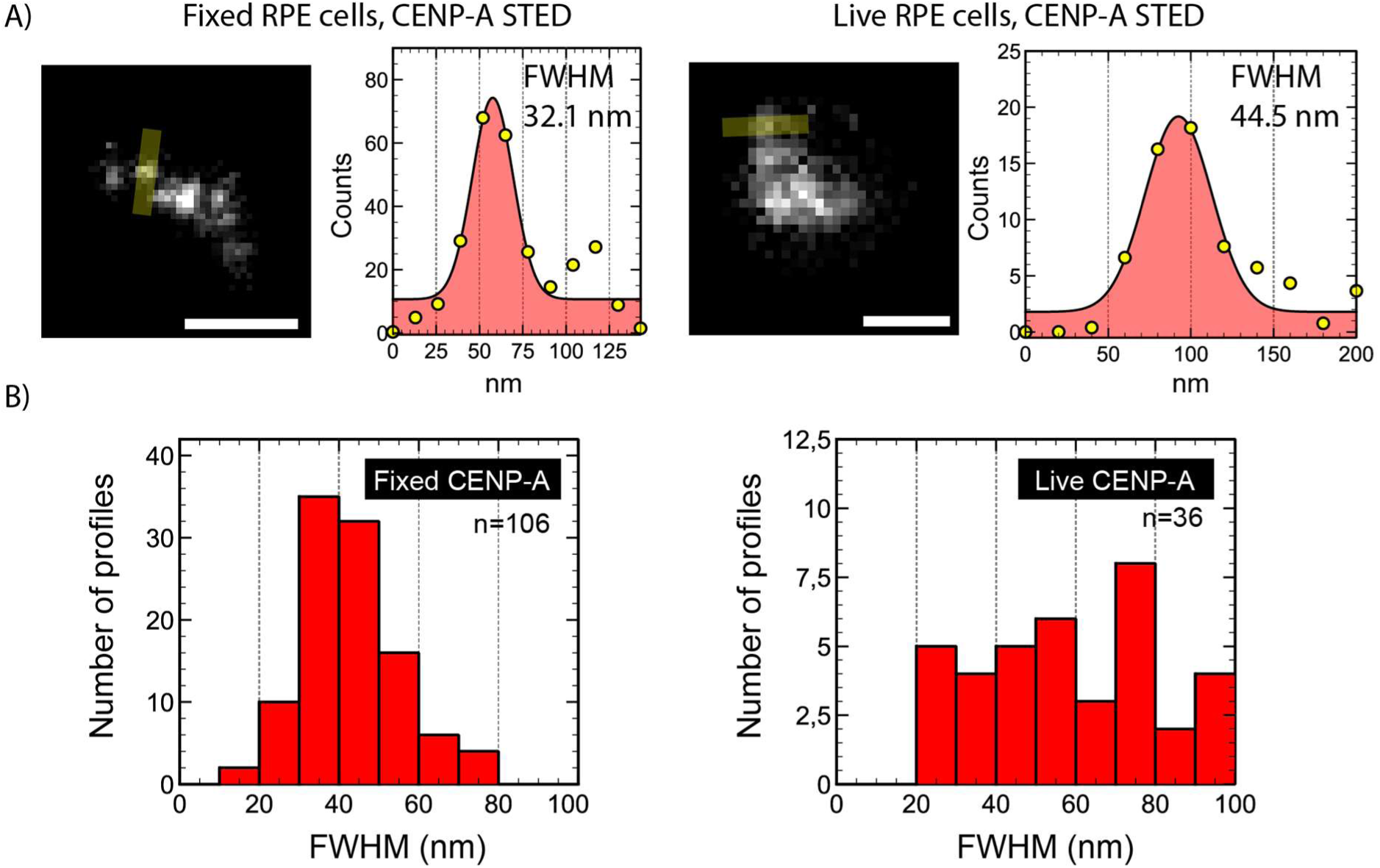
STED resolution analysis of CENP-A imaging in fixed and live RPE cells. **A)** Example line profiles (3-pixel width) drawn across CENP-A signal (yellow) with corresponding intensity plot and Gaussian fit for fixed immunolabelled CENP-A (left) and live CENP-A-Halo (right) imaging. Scale bars 200 nm. **B)** Histograms of full width at half maximum (FWHM) values obtained from Gaussian fits of 106 line profiles measured in fixed cells and 36 line profiles measured in live cells.

**Supplementary Figure S6.**
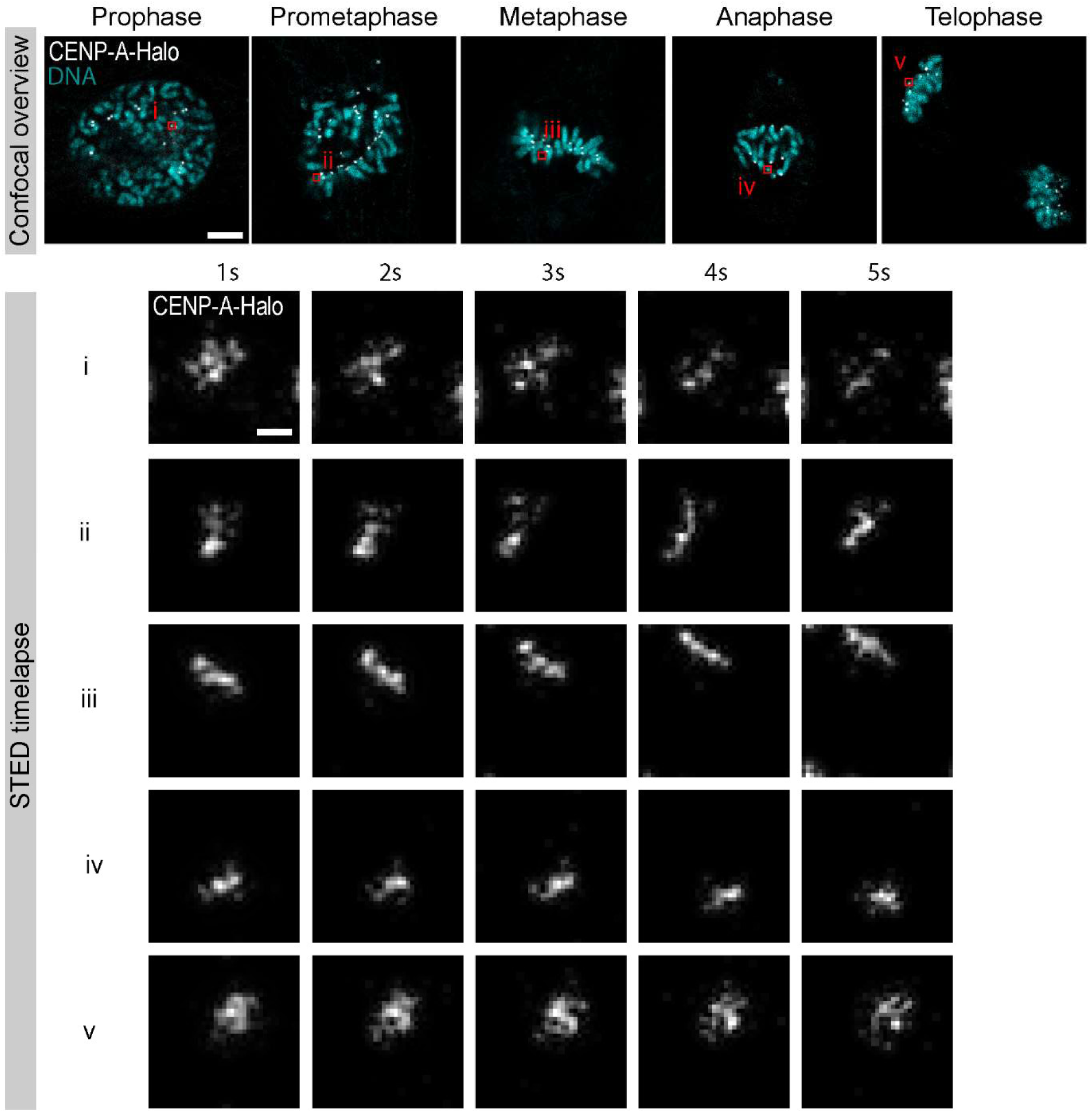
Live-cell STED time-lapse imaging of CENP-A during mitosis. hTERT RPE-1 CENP-A-Halo cells were labelled with SiR-Halo and imaged under physiological conditions. Red squares indicate the approximate region where the centromeres marked i-v resided at time of acquisition. Time-lapse STED images were acquired using DyMIN at 1 frame per second. Wiener deconvolution was applied for visualization. Each frame is intensity-scaled (min-max) to account for photobleaching. Scale bars 2 µm (confocal overview) and 200 nm (STED time-lapse).

**Supplementary Figure S7.**
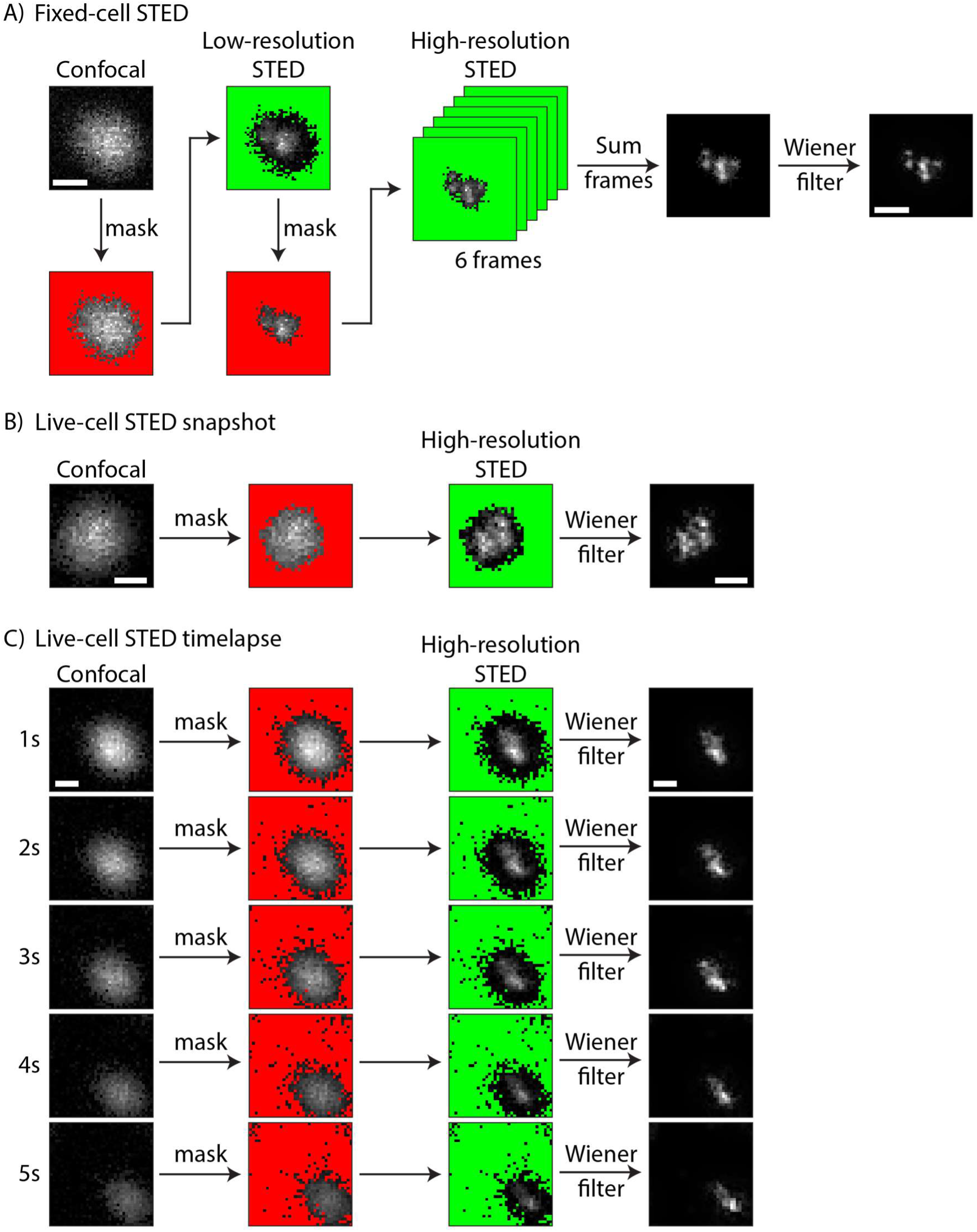
STED image acquisition and processing. STED images were made in 3 (A, fixed-cell) or 2 (B-C, live-cell) steps with intermediate masking to prevent unnecessary photobleaching. In general, a low-resolution image (confocal or low-resolution STED) is made and a mask of the background outside of the structure is obtained (red). Then, a higher-resolution image is made only of the pixels that contained signal in the low-resolution image, hence skipping high-resolution imaging of background pixels (green), which would otherwise induce photobleaching of the structure of interest. **A)** DyMIN (26) sequence for fixed-cell STED imaging: 3 imaging steps are used (confocal, low-resolution STED, and high-resolution STED) with intermediate masks to exclude background pixels. To minimize spatially inhomogeneous bleaching while obtaining maximal signal, the high-resolution STED image is acquired as 6 frames, which are summed to obtained the overall image and a Wiener filter is typically applied for improved visualization. **B)** DyMIN (26) sequence for live-cell STED snapshots: 2 imaging steps are used (confocal and high-resolution STED) and the high-resolution STED image is acquired in one frame, such that acquisition is fast and there is minimal movement during imaging. A Wiener filter is typically applied for improved visualization. **C)** DyMIN (26) sequence for live-cell STED time-lapses: 2 imaging steps are used for each frame (confocal and high-resolution STED) such that the mask is updated for every frame in case the centromere moves. A Wiener filter is typically applied for improved visualization and the intensity is rescaled per frame to account for photobleaching. Scale bars 200 nm.

